# Cannabinoid CB_2_ receptors in primary sensory neurons are implicated in CB_2_ agonist-mediated suppression of paclitaxel-induced neuropathic nociception and sexually-dimorphic sparing of morphine tolerance

**DOI:** 10.1101/2024.03.05.583426

**Authors:** Kelsey G. Guenther, Xiaoyan Lin, Zhili Xu, Alexandros Makriyannis, Julian Romero, Cecilia J. Hillard, Ken Mackie, Andrea G. Hohmann

## Abstract

Cannabinoid CB_2_ agonists show therapeutic efficacy without the unwanted side effects commonly associated with direct activation of CB_1_ receptors. The G protein-biased CB_2_ receptor agonist LY2828360 attenuates the maintenance of chemotherapy-induced neuropathic nociception in male mice and blocks the development of morphine tolerance in this model. However, the specific cell types involved in this phenomenon have never been investigated and whether this therapeutic profile is observed in female mice remains poorly understood. We used conditional deletion of CB_2_ receptors from specific cell populations to determine the population(s) mediating the anti-allodynic and morphine-sparing effects of CB_2_ agonists. Anti-allodynic effects of structurally distinct CB_2_ agonists (LY2828360 and AM1710) were present in paclitaxel-treated CB_2_^f/f^ mice of either sex. The anti-allodynic effect of the CB_2_ agonists were absent in conditional knockout (KO) mice lacking CB_2_ receptors in peripheral sensory neurons (Advillin^CRE/+^; CB_2_^f/f^) but preserved in mice lacking CB_2_ receptors in CX3CR1 expressing microglia/macrophages (CX3CR1^CRE/+^; CB_2_^f/f^). The morphine-sparing effect of LY28282360 occurred in a sexually-dimorphic manner, being present in male mice but absent in female mice of any genotype. In mice with established paclitaxel-induced neuropathy, prior LY2828360 treatment (3 mg/kg per day i.p. x 12 days) blocked the subsequent development of morphine tolerance in male CB_2_^f/f^ mice but was absent in male (or female) Advillin^CRE/+^; CB_2_^f/f^ mice. LY2828360-induced sparing of morphine tolerance was preserved in male CX3CR1^CRE/+^; CB_2_^f/f^ mice, but this effect was not observed in female CX3CR1^CRE/+^; CB_2_^f/f^ mice. Similarly, co-administration of morphine with a low dose of LY2828360 (0.1 mg/kg per day i.p. x 6 days) reversed tolerance to the anti-allodynic efficacy of morphine in paclitaxel-treated male CB_2_^f/f^ mice, but this effect was absent in female CB_2_^f/f^ mice and Advillin^CRE/+^; CB_2_^f/f^ mice of either sex. Additionally, LY2828360 (3 mg/kg per day i.p. x 8 days) delayed, but did not prevent, the development of paclitaxel-induced mechanical and cold allodynia in either CB_2_^f/f^ or CX3CR1^CRE/+^; CB_2_^f/f^ mice of either sex. Our studies reveal that CB_2_ receptors in primary sensory neurons are required for the anti-allodynic effects of CB_2_ agonists in a mouse model of paclitaxel-induced neuropathic nociception. We also find that CB_2_ agonists acting on primary sensory neurons produce a sexually-dimorphic sparing of morphine tolerance in males, but not female, paclitaxel-treated mice.

## Introduction

Morphine is an effective treatment for acute pain management, but unwanted side effects, including physical dependence, respiratory depression, and tolerance, limit its clinical use (e.g. Yekkirala et al. 2017; Coluzzi et al. 2022). Recent studies have investigated therapeutic strategies to prevent or block morphine tolerance, with a focus on co-administration of mechanistically distinct analgesics (Habibi-Asl et al. 2014; Mansouri et a. 2015; Hassanipour et al. 2016; Hosseinzadeh et al. 2016). The opioid system can functionally interact with the endogenous cannabinoid system by receptor heterodimerization or signaling cross-talk (Bushlin et al. 2010; Wolyniak et al. 2022). Indeed, co-administration of opioids with cannabinoid compounds produces synergistic antinociceptive effects in studies examining their analgesic properties (eg. Finn et al. 2004; Fischer et al. 2010; Kazantzis et al. 2016). Although activation of the cannabinoid 1 (CB_1_) receptor produces antinociception, unwanted side effects such as sedation, memory impairment and dependence also limit their clinical utility (Rahn et al. 2009; Cooper et al. 2009). By contrast, activation of the cannabinoid 2 (CB_2_) receptor alleviates various types of pathological pain without producing adverse side effects associated with CB_1_ receptor agonism (Guindon and Hohmann 2008; Deng et al. 2015). Additionally, CB_2_ receptors are involved in the antinociceptive effects of morphine under both normal and inflammatory conditions (Lim et al. 2005; Merighi et al. 2012; Desroches et al. 2014). Previous work from our group has also shown that chronic CB_2_ receptor activation can prevent or attenuate the development of morphine tolerance in various neuropathic pain models (Lin et al. 2018; Li et al. 2019; Carey et al. 2023).

LY2828360 is a potent CB_2_ receptor agonist with similar affinity for both human and rat CB_2_ receptors and low affinity for the CB_1_ receptor (Hollinshead et al. 2013). Previous studies have shown that LY2828360 suppresses behavioral hypersensitivities in models of neuropathic (Lin et al. 2018; Lin et al. 2022; Carey et al. 2023) and inflammatory (Guenther et al. 2023) pain and can prevent the development of morphine tolerance in mouse models of neuropathic pain (Lin at al. 2018; Carey et al. 2023). LY2828360 was previously evaluated in Phase 2 clinical trial for osteoarthritis pain. Although it was ineffective at treating pain in this specific population of pain patients, it was demonstrated to be safe for use in humans (Pereira et al. 2013). Given the apparent safety of LY2828360, further investigation of the potential analgesic applications for this compound is important to fully understand its range of therapeutic benefits and to identify which pain states it can best treat. Previously, our lab has identified CB_2_ receptors in peripheral sensory neurons to be necessary for antiallodynic effects of LY2828360 in both carrageenan-induced inflammatory pain (Guenther et al. 2023) and 2,3-dideoxycytidine (ddC) anti-retroviral neuropathic pain (Carey et al. 2023) using cell type specific conditional KO mice. However, the specific cell types involved in the efficacy of LY2828360 in the chemotherapy-induced neuropathic pain model or in its ability to prevent the development of tolerance to morphine’s analgesic effect in this model remain unknown.

We previously reported use of our CB_2_ EGFP reporter mouse to identify cell types (i.e. Langerhans cells, dendritic cells, keratinocytes) that were dynamically regulated by challenge with the chemotherapeutic agent paclitaxel. Here, we generated conditional knockout (KO) mouse lines with deletion of CB_2_ receptor from peripheral sensory neurons (Advillin^CRE/+^; CB_2_^f/f^ mice) or microglia/macrophages expressing the protein coding gene C-X3-C Motif Chemokine Receptor 1 (CX3CR1^CRE/+^; CB_2_^f/f^ mice) to examine the contribution of these cell types to the development and maintenance of chemotherapy-induced neuropathic pain and the therapeutic effects of CB_2_ agonists. We also compared effects of LY2828360 with those of AM1710, a CB_2_ receptor agonist that differs in its ability to penetrate the central nervous system (CNS; Rahn et al. 2011). Additionally, we investigated the ability of LY2828360 to prevent or reverse morphine tolerance in neuropathic pain in CB_2_^f/f^ and our conditional KO mice. Lastly, we examined the impact of prophylactic treatment with LY2828360 in paclitaxel-treated CX3CR1^CRE/+^; CB_2_^f/f^ mice. Our studies document a previously unrecognized role for CB_2_ receptors in primary sensory neurons in anti-allodynic effects of CB_2_ agonists in chemotherapy-induced neuropathic nociception and document a role for these receptors in a sex-specific sparing of morphine tolerance that we observe in paclitaxel-treated male, but not female, CB_2_^f/f^ mice.

## Methods

### Subjects

Adult male C57BL6/J mice were purchased from Jackson labs (Bar Harbor, ME) for qRT-PCR experiment. All conditional KO mice were bred on a C57BL6/J background at Indiana University and included adult male and female CB_2_^f/f^, Advillin^CRE/+^; CB_2_^f/f^ (deletion of CB_2_ from peripheral sensory neurons) and CX3CR1^CRE/+^; CB_2_^f/f^ mice (deletion of CB_2_ from microglia/macrophages). To conditionally delete CB_2_ from the desired cells, female mice carrying two alleles of the floxed CB_2_ gene (CB_2_^f/f^; López et al. 2018) were bred with male mice carrying two floxed CB_2_ alleles and one allele of Cre-recombinase in either the CX3CR1 (Yona et al. 2013) or Advillin (Zhou et al. 2010; da Silva et al. 2011). All mice were housed in a temperature-controlled colony room on a 24-hr light-dark cycle. Behavioural testing occurred during the light phase of the cycle. Mice were maintained on ad-libitum food and water. All experiments were approved by the Indiana University Animal Care and Use Committee.

### Drugs

Paclitaxel (Tecoland Corporation, Irvine, CA) was dissolved in a cremophor-based vehicle made of Cremophor EL (Sigma-Aldrich, St. Louis, MO), ethanol (Sigma-Aldrich) and 0.9% saline (Aqualite System; Hospira, Inc., Lake Forest, IL) at a 1:1:18 ratio as previously published (Deng et al., 2015). LY2828360 (8-(2-chlorophenyl)-2-methyl-6-(4-methylpiperazin-1-yl)-9-(tetrahydro-2H-pyran-4-yl)-9H-purine) was synthesized by Sai Biotech (Mumbai, India; purity >98%). AM1710 was synthesized by the lab of Alexandros Makriyannis (Northeastern University, Boston, MA). Morphine (Sigma-Aldrich), AM1710 and LY2828360 were dissolved in a vehicle containing 10% dimethyl sulfoxide (DMSO; Sigma-Aldrich), and the remaining 90% consisted of emulphor (Alkamuls EL-620; Rhodia, Cranbury, NJ), ethanol and saline at a 1:1:18 ratio. Naloxone (Sigma-Aldrich) was dissolved in saline. All compounds were administered intraperitoneally (i.p.) at a volume of 10 ml/kg.

### Tissue collection and quantitative real-time polymerase chain reaction (qRT-PCR)

In Experiment 1, real-time (RT) quantitative PCR (qPCR) was used to examine the distribution of GFP mRNA (used as a surrogate marker for CB_2_ mRNA in CB2f/f mice) and CB_2_ mRNA in the dorsal root ganglia (DRG), lumbar spinal cord, paw skin and spleen of male and female naïve wild type (C57BL/6J), CB_2_^f/f^ and Advillin^CRE/+^; CB_2_^f/f^ mice (n =5-13 per group). Mice were perfused with Hank’s balanced Salt solution (HBBS), then DRG, spinal cord, paw skin and spleen were collected and flash frozen in liquid nitrogen then stored at – 80 °C until use. DRG and paw skin RNA was extracted using a RLT buffer:2-mercapto ethanol mixture in a 100:1 ratio/RNeasy (Qiagen) RNA mini kit according to the manufacturer’s instructions and spinal cord and spleen RNA was extracted using TRizol (Invitrogen)/RNeasy RNA mini kit. Quantification of total RNA was assessed using NanoDrop 2000C UV-Vis spectrophotometer at 260 nm.

One-step RT-qPCR was performed using the Luna universal One-Step RT-qPCR kit (New England Biolabs) in a total volume of 10 µl and a template concentration of 50 ng/µl according to manufacturer’s recommendation. Thermal cycle conditions were 55 °C for 10 min, 95 °C for 1 min, followed by 40 cycles of 95 °C for 10 sec and 60 °C for 1 min. A melting curve analysis was performed at 95 °C for 15 sec, 60 °C for 1 min, 95 °C for 15 sec following every run to ensure a single amplified product for each reaction. All reactions were performed in duplicate in 384-well reaction plates (Thermo Fisher Scientific). The mRNA expression levels were quantified by cycle threshold (Ct), where Ct levels are inversely proportional to the amount of target nucleic acid in the sample. Relative gene expression for CB2 and GFP were normalized to the housekeeping gene GAPDH and calculated using the 2^-ΔΔCt^ method (Livak and Schmittgen, 2001). Sequences for qRT-PCR primers were: CB_2_ sense: 5’-CTCGGTTACAGAAACAGAGGCTGATGTG-3’; CB_2_ anti-sense: 5’-

TCTCTCTTCGAGGGAGTGAACTGAACG-3’; GFP sense: 5’-ACATGGTCCTGCTGGAGTTCGTGAC-3’; GFP anti-sense: 5’-CTCTTCGAGGGAGTGAACTGAACGG-3’; GAPDH sense: 5’-GGGAAGCTCACTGGCATGGC-3’; anti-sense: 5’-GGTCCACCACCCTGTTGCT-3’.

### Behavioral hypersensitivity testing

Paw withdrawal thresholds to mechanical stimulation were assessed using an electronic von Frey anesthesiometer (IITC, Woodland Hills, CA) as described in our previously published work (Lin et al. 2018, 2022). Animals were placed on an elevated testing table with a stainless-steel wire mesh platform (0.6 x 0.6 cm gaps in the wire mesh) underneath clear Plexiglass chambers (10.5 x 9 x 7 cm) for at least 20 minutes prior to testing. Following this habituation period, a force was applied to the midplantar region of the hind paw until a paw withdrawal response was observed. Mechanical stimulation was terminated when the animal withdrew its paw, and the value of the maximum applied force was recorded in grams. Each paw was measured twice at every time point, and values were averaged for each animal.

Sensitivity to cold was assessed immediately following testing for mechanical paw withdrawal thresholds by applying a droplet of acetone (Sigma-Aldrich; approximately 5-6 μL) from the end of a blunt one C.C. syringe hub onto the midplantar region of the hind paw. Total time the mouse spent attending to the acetone-stimulated paw (ie. elevation, shaking or licking) was recorded over a 1-minute period. Each paw was measured three times at every time point, and values were averaged for each animal.

### General in vivo experimental protocol

In all studies, the experimenter was blind to the treatment conditions and mice were randomly assigned to experimental conditions. Paclitaxel (4 mg/kg i.p.) was administered four times on alternate days (cumulative dose of 16 mg/kg) to induce neuropathic pain as described previously (Deng et al., 2015). Control mice received an equal volume of cremophor vehicle. Paclitaxel-induced peripheral neuropathy was established prior to pharmacological manipulation, except in the prophylactic study. Mechanical and cold allodynia were measured 30 minutes after drug administration on testing days.

In Experiment 2, we evaluated whether CB_2_^f/f^ and Advillin^CRE/+^; CB_2_^f/f^ mice develop mechanical and cold hypersensitivities in response to treatment with paclitaxel or its vehicle (n = 4-7 per group).

In Experiment 3, we compared the antiallodynic efficacy of two structurally distinct CB_2_ agonists that differ in their ability to penetrate the central nervous system (CNS) using both CB_2_^f/f^ and Advillin^CRE/+^; CB_2_^f/f^ mice (n = 5-6 per group). Male mice were given an i.p. injection of either AM1710, a CB_2_ agonist with limited CNS penetration (Rahn et al., 2011), or LY2828360, a CNS-penetrant CB_2_ agonist (Hollinshead et al. 2013), in a dose-response experiment. Escalating doses of AM1710 (1, 3, 5 and 10 mg/kg) or LY2828360 (0.1, 0.3, 1 and 3 mg/kg) were examined for antiallodynic efficacy using a within-subject escalating dosing paradigm. Successive doses were separated by 2 to 4 days.

In Experiment 4, we examined the antiallodynic efficacy of chronic LY2828360 administration (3 mg/kg per day i.p. x 12 days) or vehicle administered once daily for 12 consecutive days (i.e. phase I) in both male and female CB_2_^f/f^ and Advillin^CRE/+^; CB_2_^f/f^ mice. This treatment was followed approximately 4 days later by a second phase of once daily dosing for 12 consecutive days (i.e. phase II) of chronic morphine (10 mg/kg per day i.p. x 12 days) or vehicle in the same animals (n = 8-9 per group). Responsiveness to mechanical and cold stimulation were measured on days 1, 4, 8 and 12 during phase I and on days 16, 19, 23 and 27 during phase II (phase II began on day 16).

In Experiment 5, we examined the antiallodynic efficacy of low dose LY2828360 or vehicle in both male and female CB_2_^f/f^ and Advillin^CRE/+^; CB_2_^f/f^ mice that had previously developed tolerance to morphine (n = 6-8 per group). To induce morphine tolerance, mice received repeated daily injections of morphine in phase I (10 mg/kg per day i.p x 6 days), followed by low dose LY2828360 (0.1 mg/kg per day i.p. x 6 days) or vehicle co-administered with morphine (10 mg/kg per day i.p x 6 days) in phase II. Responsiveness to mechanical and cold stimulation were measured on days 1, 3 and 6 during phase I and on days 7, 9 and 12 during phase II (phase II began on day 7).

In Experiment 6, we evaluated the antiallodynic efficacy of LY2828360 in both CB_2_^f/f^ and CX3CR1^CRE/+^; CB_2_^f/f^ mice as described in experiment 3 (n = 6 for cremophor groups (female only); n = 16 for paxlitaxel groups (mixed sex)).

In Experiment 7, we examined the antiallodynic efficacy of chronic LY2828360 or vehicle administered during phase I in both male and female CB_2_^f/f^ and CX3CR1^CRE/+^; CB_2_^f/f^ mice, followed by the assessment of the antiallodynic efficacy of chronic morphine administration during phase II in the same animals, as described in Experiment 4 (n = 6-8 per group).

In Experiment 8, we examined the ability of LY2828360 (3 mg/kg per day i.p. x 8) to prevent the development of paclitaxel-induced mechanical allodynia. Paclitaxel (4 mg/kg i.p.) was injected on days 0, 2, 4 and 6, and LY2828360 was injected on days 0-7. LY2828360 was injected 1-hr prior to paclitaxel on days when both drugs were administered. Responsiveness to mechanical and cold stimulation were measured on days 0, 4, 7, 11, 14 and 18, with behavioral testing performed prior to daily injections.

### Evaluation of opioid receptor-mediated withdrawal symptoms

Mice that received morphine (10 mg/kg per day i.p.) following LY2828360 administration (3 mg/kg per day i.p.) or a combination of morphine and LY2828360 (10 mg/kg per day i.p. morphine co-administered with 0.1 mg/kg per day i.p. LY2828360) in the two-phase studies were challenged with naloxone (5 mg/kg i.p.) to induce opioid withdrawal. Specifically, mice used in Experiments 4, 5, and 7 were challenged with naloxone to precipitate opioid withdrawal beginning 30 minutes after the last injection of the test drug. Mice were video recorded, and the number of naloxone-precipitated withdrawal jumps was scored over a 30-minute observation period.

## Results

### Experiment 1. Both GFP and CB_2_ mRNAs are reduced in the dorsal root ganglia of Advillin^CRE/+^; CB_2_^f/f^ relative to CB_2_^f/f^ mice

Advillin^CRE/+^; CB_2_^f/f^ mice showed reduced CB_2_ mRNA in the DRG compared to both CB_2_^f/f^ (**DRG:** *p* = 0.0002; **SC:** *p* = 0.0780; **PS:** *p* = 0.8159; **spleen:** *p* = 0.0726) and wildtype (**DRG:** *p* < 0.0001; **SC:** *p* = 0.5148; **PS:** *p* = 0.6373; **spleen:** *p* = 0.0955) mice whereas no such changes were observed in lumbar spinal cord (SC), paw skin (PS) or spleen. Overall, CB_2_ mRNA expression levels were similar between CB_2_^f/f^ and wildtype mice with one exception; a modest but reliable increase in CB_2_ mRNA expression levels was observed in lumbar spinal cord of CB_2_^f/f^ mice relative to wild type mice (*p* = 0.0447). Thus, GFP mRNA expression levels tracked CB_2_ mRNA expression levels in CB_2_^f/f^ mice in all tissues and were elevated relative to wildtype (background) levels that, as expected, did not express GFP mRNA (**DRG:** *p* < 0.0001; **SC:** *p* < 0.0001; **PS:** *p* < 0.0001; **spleen:** *p* < 0.0001). In DRG, GFP mRNA expression levels were lower in advillin^CRE/+^; CB_2_^f/f^ mice compared to CB_2_^f/f^ mice (*p* < 0.0001), but not in other tissues (**Fig. 1**).

**Fig. 1.**
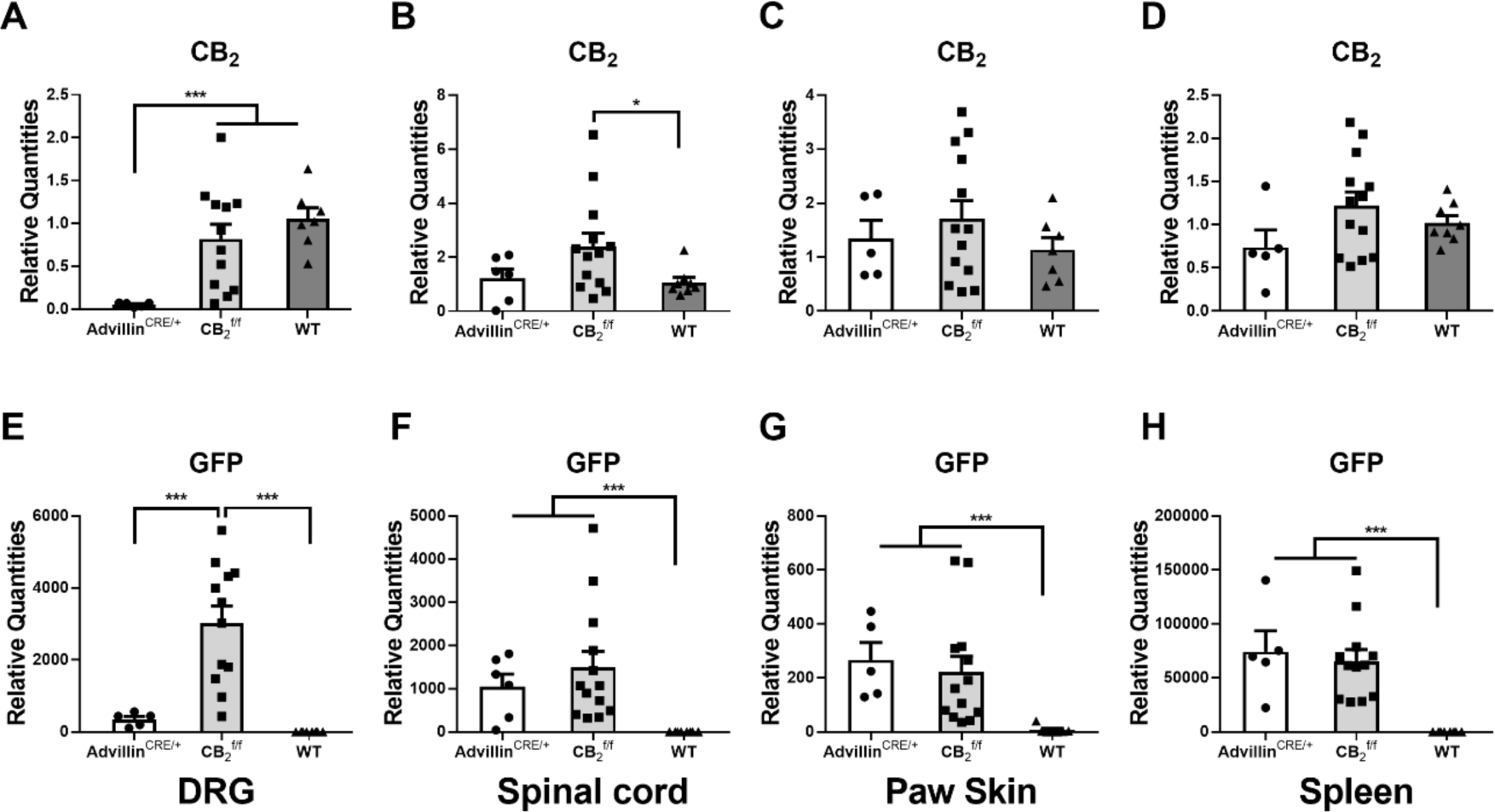
Comparison of CB_2_ mRNA (A, B, C, D) and GFP mRNA (E, F, G, H) expression in DRG (A, E), spinal cord (B, F), paw skin (C, G) and spleen (D, H) of male and female Advillin^CRE/+^; CB_2_^f/f^, CB_2_^f/f^ and wild type (WT) mice. Advillin^CRE/+^; CB_2_^f/f^ mice showed lower levels of CB_2_ mRNA compared to both CB_2_^f/f^ and WT mice and lower levels of GFP compared to CB_2_^f/f^ mice. WT mice did not show any GFP mRNA expression. Mean (±SEM) 2^-ΔΔCT^ values for experiment 1 (n = 5-13 per group). Asterisk indicates a group difference; \**p* < 0.05, \*\*\**p* < 0.001.

### Experiment 2. The chemotherapy agent paclitaxel induced hypersensitivity to mechanical and cold stimulation in both CB_2_^f/f^ and Advillin^CRE/+^; CB_2_^f/f^ mice

Paclitaxel decreased mechanical paw withdrawal thresholds and increased cold sensitivity as a function of time in both CB_2_^f/f^ and Advillin^CRE/+^; CB_2_^f/f^ mice. (**Mechanical:** Time: F_4,64_ = 24.42, *p* < 0.0001; Group: F_3,16_ = 77.53, *p* < 0.0001; Interaction: F_12,64_ = 6.71, *p* < 0.0001; **Cold:** Time: F_4,64_ = 21.76, *p* < 0.0001; Group: F_3,16_ = 7.78, *p* =0.0020; Interaction: F_12,64_ = 5.696, *p* < 0.0001). Sidak’s multiple comparisons test revealed decreased mechanical withdrawal threshold (*p* < 0.0001 on day 4, 7, 10 and 14), and increased cold sensitivity (*p* < 0.0043 on day 7, 10 and 14), following paclitaxel (Pac) treatment compared to cremophor (crem) vehicle treatment, in both CB_2_^f/f^ (**Fig. 2A**) and Advillin^CRE/+^; CB_2_^f/f^ (**Fig. 2B**) mice. Responses to mechanical and cold stimulation did not differ between CB_2_^f/f^ and Advillin^CRE/+^; CB_2_^f/f^ mice in either the paclitaxel or crem vehicle-treated groups.

**Fig. 2.**
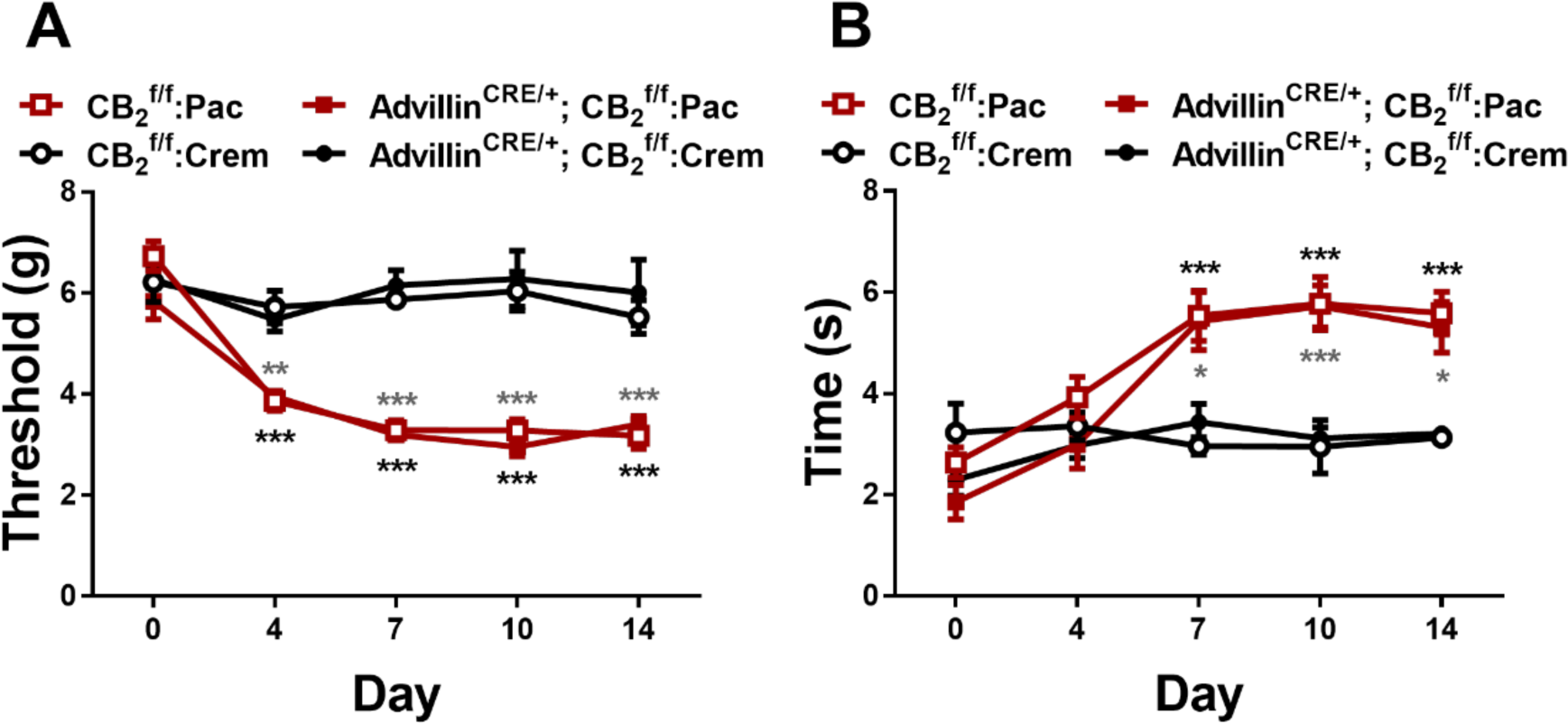
Mechanical (A) and cold (B) allodynia develop normally in CB_2_^f/f^ and Advillin^CRE/+^; CB_2_^f/f^ mice following paclitaxel treatment (n = 4-7 per group). Data expressed as mean (±SEM), \**p* < 0.05, \*\**p* < 0.005, \*\*\**p* < 0.001. Gray asterisk indicate difference between CB_2_^f/f^ groups; black asterisk indicate difference between Advillin^CRE/+^; CB_2_^f/f^ groups.

### Experiment 3. The CB_2_ agonists AM1710 and LY2828360 dose dependently reduced mechanical and cold allodynia in CB_2_^f/f^, but not Advillin^CRE/+^; CB_2_^f/f^ mice

LY2828360 increased mechanical paw withdrawal thresholds and decreased cold sensitivity as a function of dose and genotype in paclitaxel-treated mice (**Mechanical:** Dose: F_4,40_ = 24.21, *p* < 0.0001; Genotype: F_1,10_ = 74.71, *p* < 0.0001; Interaction: F_4,40_ = 21.81, *p* < 0.0001; **Cold:** Dose: F_4,40_ = 6.441, *p* = 0.0004; Genotype: F_1,10_ = 10.05, *p* = 0.0100; Interaction: F_4,40_ = 11.13, *p* < 0.0001). Sidak’s multiple comparison post hoc test revealed that LY2828360 increased mechanical withdrawal thresholds at doses of 0.1 (*p* < 0.0301), 0.3 (*p* < 0.0239), 1 and 3 mg/kg (*p* < 0.0001; **Fig. 3A**) in CB_2_^f/f^ mice but not Advillin^CRE/+^; CB_2_^f/f^ mice. LY2828360 also decreased cold sensitivity at doses of 1 and 3 mg/kg (*p* < 0.0001; **Fig. 3B**) in CB_2_^f/f^ mice but not Advillin^CRE/+^; CB_2_^f/f^ mice. LY2828360 had no effect on mechanical paw withdrawal thresholds (**Fig. 3C**) in either CB_2_^f/f^ or Advillin^CRE/+^; CB_2_^f/f^ mice treated with cremophor-based vehicle in lieu of paclitaxel (**Mechanical:** Dose: F_4,32_ = 0.3443, *p* = 0.8460; Genotype: F_1,8_ = 0.007119, *p* = 0.9348; Interaction: F_4,32_ = 0.8399, *p* = 0.5101). Cold responsiveness changed as a function of dose and was slightly elevated in CB_2_^f/f^ relative to Advillin^CRE/+^; CB_2_^f/f^ mice treated with cremophor-based vehicle, but the interaction was not significant (**Cold:** Dose: F_4,32_ = 3.625, *p* = 0.0152; Genotype: F_1,8_ = 5.8, *p* = 0.0426; Interaction: F_4,32_ = 0.3422, *p* = 0.8474; **Fig. 3D**).

**Fig. 3.**
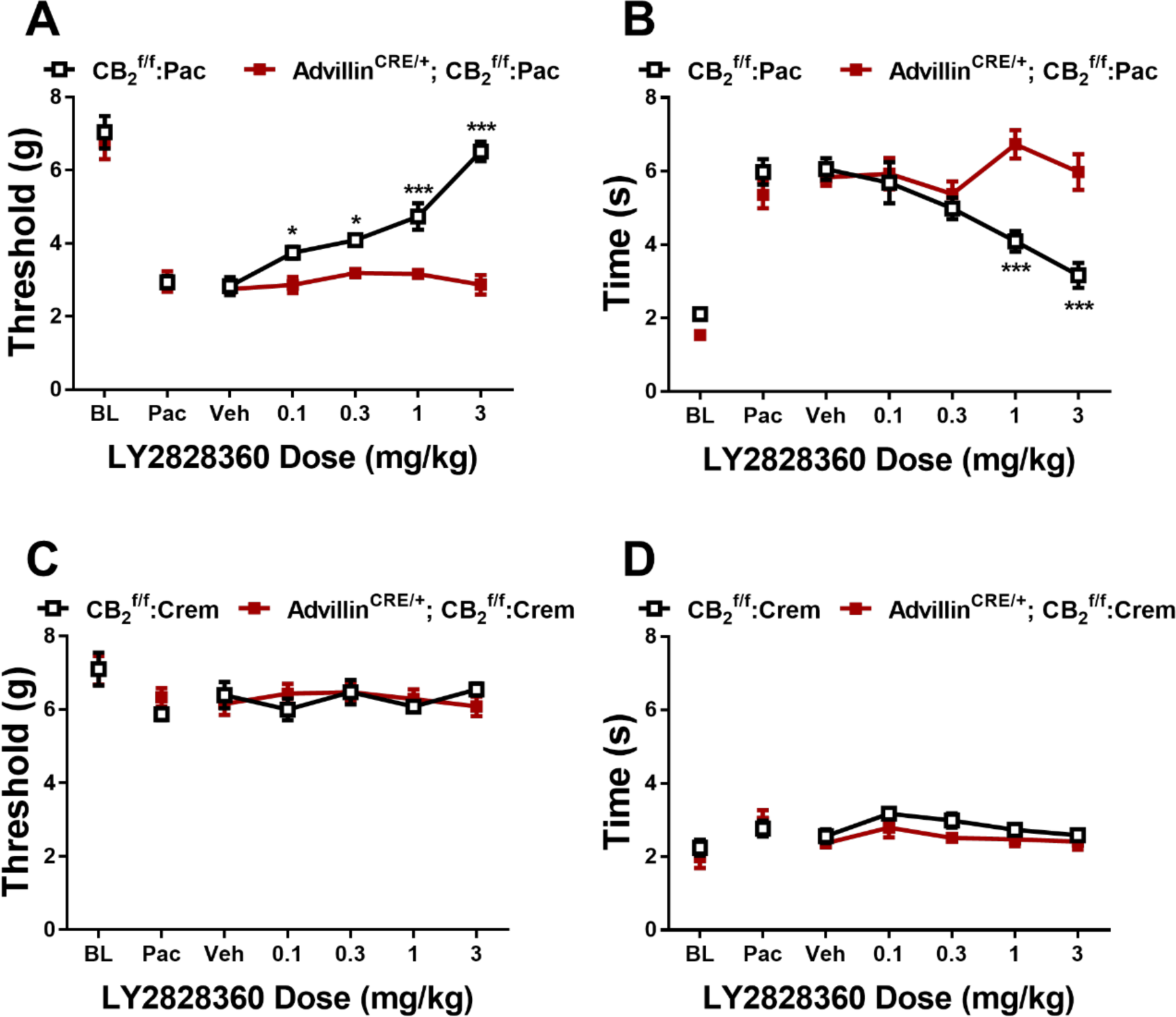
LY2828360 dose dependently alleviated paclitaxel evoked mechanical (A) and cold (B) allodynia in CB_2_^f/f^, but not Advillin^CRE/+^; CB_2_^f/f^ mice. LY2828360 did not alter mechanical (C) or cold (D) sensitivity in CB_2_^f/f^ or Advillin^CRE/+^; CB_2_^f/f^ control mice treated with cremophor vehicle (n = 5-6 per group). Data expressed as mean (±SEM), \**p* < 0.05, \*\*\**p* < 0.001.

In paclitaxel-treated mice, AM1710 increased mechanical paw withdrawal thresholds and decreased cold sensitivity as a function of dose and genotype (**Mechanical:** Dose: F_4,40_ = 13.29, *p* < 0.0001; Genotype: F_1,10_ = 24.48, *p* = 0.0006; Interaction: F_4,40_ = 6.385, *p* = 0.0005; **Cold:** Dose: F_4,40_ = 3.782, *p* = 0.0106; Genotype: F_1,10_ = 5.3, *p* = 0.0106; Interaction: F_4,40_ = 6.386, *p* = 0.0005). Sidak’s multiple comparisons test revealed that AM1710 increased mechanical withdrawal thresholds at doses of 5 and 10 mg/kg (*p* < 0.0001; **Fig. 4A**) and decreased cold sensitivity at doses of 5 (*p* = 0.0080) and 10 mg/kg (*p* = 0.0007; **Fig. 4B**) in CB_2_^f/f^ but not Advillin^CRE/+^; CB_2_^f/f^ mice. AM1710 did not alter mechanical or cold responsiveness at doses of 1 and 3 mg/kg (*p* > 0.1331) in either genotype. AM1710 did not alter mechanical (**Fig. 4C**) or cold (**Fig. 4D**) responsiveness in either CB_2_^f/f^ or Advillin^CRE/+^; CB_2_^f/f^ mice treated with cremophor-based vehicle (**Mechanical:** Dose: F_4,32_ = 1.486, *p* = 0.2297; Genotype: F_1,8_ = 9.052e-005, *p* = 0.9926; Interaction: F_4,32_ = 0.338, *p* = 0.8503; **Cold:** Dose: F_4,32_ = 5922, *p* = 0.6708; Genotype: F_1,8_ = 5.8, *p* = 0.8780; Interaction: F_4,32_ = 0.6736, *p* = 0.6151).

**Fig. 4.**
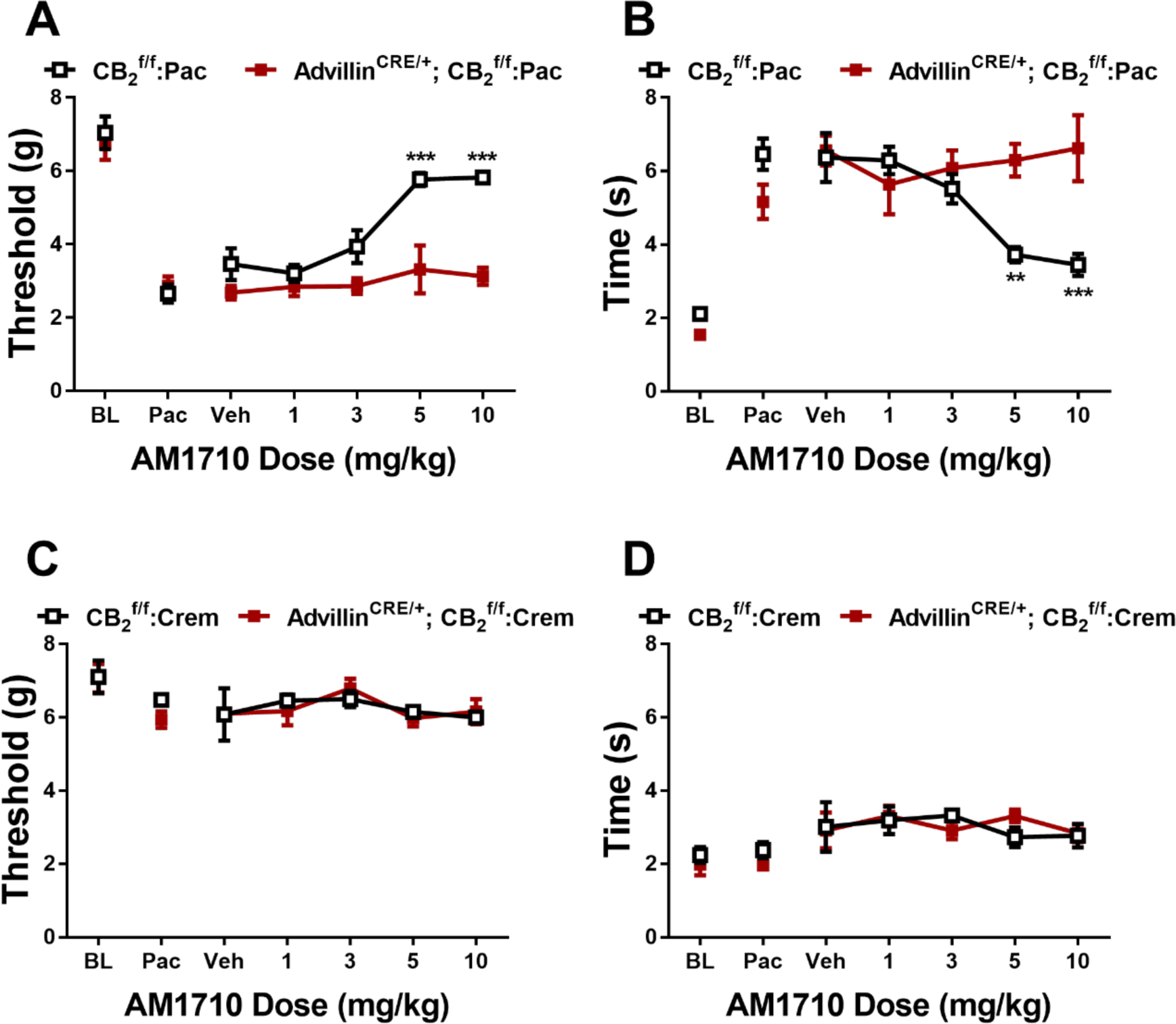
AM1710 dose dependently alleviated paclitaxel-evoked sensitization to mechanical (A) and cold (B) stimulation in CB_2_^f/f^, but not Advillin^CRE/+^; CB_2_^f/f^ mice. AM1710 did not alter mechanical (C) or cold (D) sensitivity in CB_2_^f/f^ or Advillin^CRE/+^; CB_2_^f/f^ control mice treated with cremophor vehicle (n = 5-6 per group). Data expressed as mean (±SEM), \*\**p* < 0.005, \*\*\**p* < 0.001.

### Experiment 4. Prior chronic dosing with LY2828360 blocked the development of morphine tolerance in male but not female CB_2_^f/f^ mice and this effect was absent in Advillin^CRE/+^; CB_2_^f/f^ mice

To examine the effects of LY2828360 treatment on the development of morphine tolerance in Advillin^CRE/+^; CB_2_^f/f^ mice, a two-phase pharmacological manipulation was used during the maintenance of neuropathic pain (**Fig. 5A**). Phase I treatment with LY2828360 (3 mg/kg per day i.p. x 12 days) increased mechanical withdrawal thresholds (**Fig. 5B,D**) and decreased cold sensitivity (**Fig. 5C,E**) in CB_2_^f/f^ but not Advillin^CRE/+^; CB_2_^f/f^ mice. This effect was observed in both male (**Mechanical:** Time: F_5,75_ = 35.66, *p* < 0.0001; Genotype: F_1,15_ = 191, *p* < 0.0001; Interaction: F_5,75_ = 27.37, *p* < 0.0001; **Cold:** Time: F_5,75_ = 73.97, *p* < 0.0001; Genotype: F_1,15_ = 144.6, *p* < 0.0001; Interaction: F_5,75_ = 39.65, *p* < 0.0001) and female (**Mechanical:** Time: F_5,70_ = 32.81, *p* < 0.0001; Genotype: F_1,14_ = 62.4, *p* < 0.0001; Interaction: F_5,70_ = 15.67, *p* < 0.0001; **Cold:** Time: F_5,70_ = 100.3, *p* < 0.0001; Genotype: F_1,14_ = 75.06, *p* < 0.0001; Interaction: F_5,70_ = 43.11, *p* < 0.0001) mice; Sidak’s multiple comparisons test revealed that LY2828360 increased mechanical paw withdrawal thresholds and decreased cold responsiveness in CB_2_^f/f^ mice compared to Advillin^CRE/+^; CB_2_^f/f^ mice on all phase I days tested in paclitaxel-treated mice of both sexes (*p* < 0.0001). Phase II treatment with morphine (10 mg/kg per day i.p. x 12 days) increased mechanical paw withdrawal thresholds (**Fig. 5B,D**) and decreased cold sensitivity (**Fig. 5C,E**) without the development of tolerance in male CB_2_^f/f^ but not male Advillin^CRE/+^; CB_2_^f/f^ mice (**Mechanical:** Time: F_6,90_ = 54.69, *p* < 0.0001; Genotype: F_1,15_ = 104.8, *p* < 0.0001; Interaction: F_6,90_ = 21.81, *p* < 0.0001; **Cold:** Time: F_6,90_ = 90.56, *p* < 0.0001; Genotype: F_1,15_ = 72.37, *p* < 0.0001; Interaction: F_6,90_ = 42.22, *p* < 0.0001). This sparing of morphine tolerance was absent in female mice of either genotype (**Mechanical:** Time: F_6,84_ = 112.5, *p* < 0.0001; Genotype: F_1,14_ = 0.007455, *p* = 0.9324; Interaction: F_6,84_ = 0.7039, *p* = 0.6473; **Cold:** Time: F_6,84_ = 386.8, *p* < 0.0001; Genotype: F_1,14_ = 2.87, *p* = 0.1124; Interaction: F_6,84_ = 4.923, *p* = 0.0002); Sidak’s multiple comparisons test revealed that acute treatment with morphine in Phase II (day 16) increased mechanical paw withdrawal thresholds and decreased cold sensitivity compared to day 15 post-paclitaxel baseline responding in both sexes regardless of genotype (*p* < 0.0001 for all comparisons). Morphine tolerance developed in male Advillin^CRE/+^; CB_2_^f/f^ and female mice of both genotypes at subsequent time points (*p* < 0.0001 on days 19-27 for all timepoints compared to day 16), whereas anti-allodynic effects of morphine were preserved in paclitaxel-treated CB_2_^f/f^ male mice for the duration of phase 2 (*p* < 0.0001 on days 19-27 compared to male Advillin^CRE/+^; CB_2_^f/f^ mice).

**Fig. 5.**
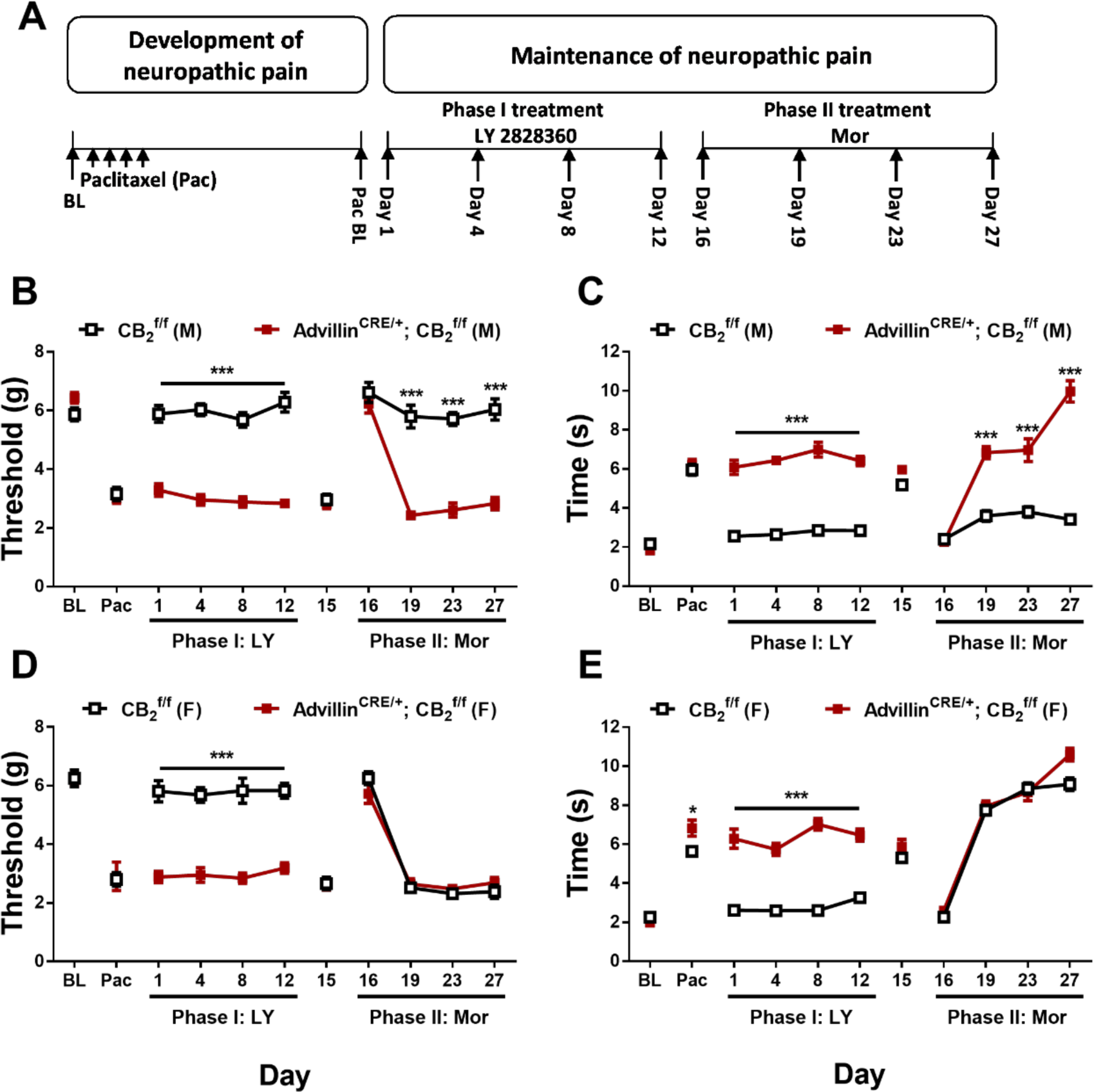
History of chronic LY2828360 treatment blocked the development of morphine tolerance in male, but not female, CB_2_^f/f^ mice, whereas morphine tolerance developed in Advillin^CRE/+^; CB_2_^f/f^ mice of either sex. (A) Schematic shows the injection schedule used to evaluate the two-phase treatment during the maintenance phase of paclitaxel-induced neuropathic pain. Chronic LY2828360 treatment (3 mg/kg per day i.p. x 12 days) alleviated paclitaxel evoked mechanical (B) and cold (C) allodynia in male CB_2_^f/f^, but not Advillin^CRE/+^; CB_2_^f/f^ mice and blocked the development of tolerance to the anti-allodynic effects of morphine (10 mg/kg per day i.p. x 12 days). LY2828360 had no effect on mechanical (D) or cold (E) allodynia in female CB_2_^f/f^ or Advillin^CRE/+^; CB_2_^f/f^ mice and did not block the development of tolerance to the anti-allodynic effects of morphine (n = 8-9 per group). Data expressed as mean (±SEM), \*\*\**p* < 0.001.

In paclitaxel-treated CB_2_^f/f^ and Advillin^CRE/+^; CB_2_^f/f^ mice with a history of chronic treatment with LY282860 and morphine, naloxone challenge produced characteristic jumping behavior that was higher in females compared to males but did not differ as a function of genotype and the interaction was not significant (Sex: F_1,29_ = 6.69, *p* = 0.0150; Genotype: F_1,29_ = 1.591, *p* = 0.2172; Interaction: F_1,29_ = 0.4572, *p* = 0.5043; **Fig. 6**).

**Fig. 6.**
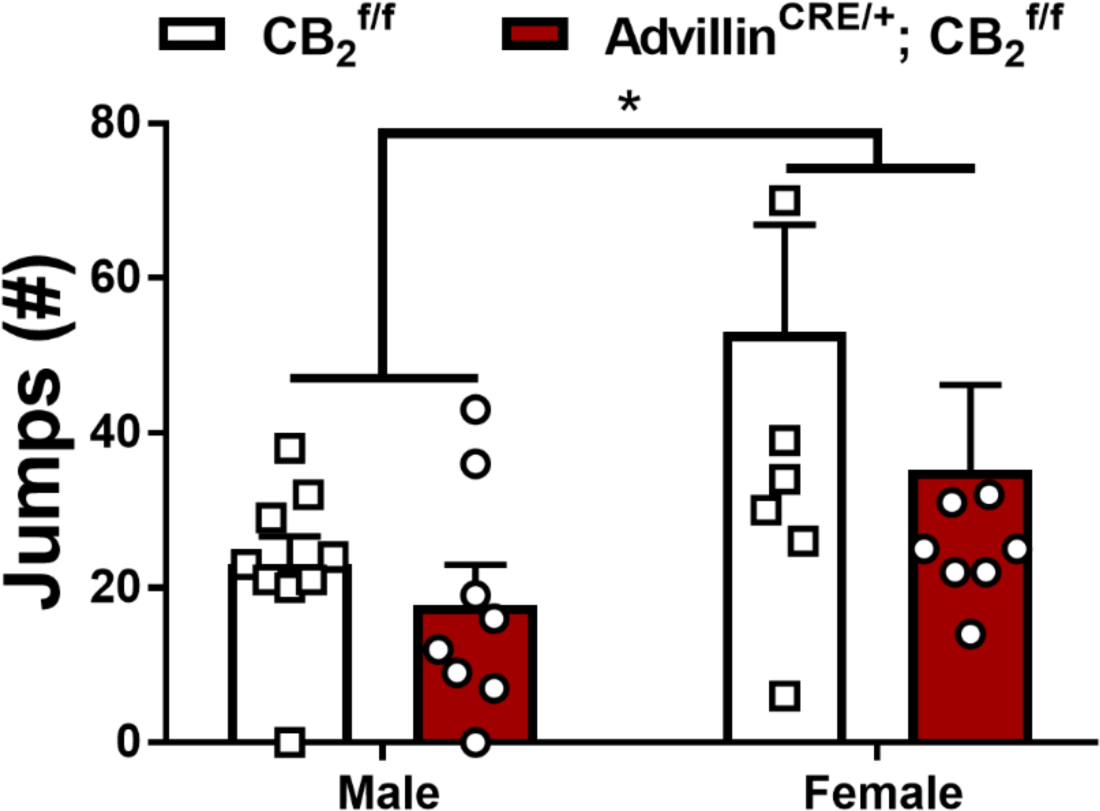
Impact of a history of chronic LY2828360 (3 mg/kg per day i.p. x 12 days) on naloxone-precipitated morphine withdrawal in male and female CB_2_^f/f^ and Advillin^CRE/+^; CB_2_^f/f^ mice. Following chronic treatment with morphine (10 mg/kg per day i.p. x 12 days) male mice show fewer withdrawal jumps compared to female mice regardless of genotype (n = 8-9 per group). Data expressed as mean (±SEM), \**p* < 0.05.

### Experiment 5. LY2828360 reversed morphine tolerance in male, but not female, CB_2_^f/f^ mice when co-administered with morphine and did not block tolerance in either male or female Advillin^CRE/+^; CB_2_^f/f^ mice

We evaluated whether a low dose of LY2828360, administered concurrently with morphine during the maintenance of neuropathic pain would reverse morphine tolerance after it already developed (**Fig. 7A**). Phase I treatment with morphine (10 mg/kg per day i.p. x 6 days) increased mechanical withdrawal thresholds (**Fig. 7B,D**) and decreased cold sensitivity (**Fig. 7C,E**) in both male (**Mechanical:** Time: F_5,120_ = 206.5, *p* < 0.0001; Group: F_3,24_ = 1.374, *p* = 0.2748; Interaction: F_15,120_ = 0.9599, *p* = 0.5015; **Cold:** Time: F_5,120_ = 302.9, *p* < 0.0001; Group: F_3,24_ = 0.5196, *p* = 0.6728; Interaction: F_15,120_ = 0.9665, *p* = 0.4945) and female (**Mechanical:** Time: F_5,125_ = 187.5, *p* < 0.0001; Group: F_3,25_ = 0.2581, *p* = 0.8549; Interaction: F_15,125_ = 1.575, *p* = 0.0899; **Cold:** Time: F_5,120_ = 392.8, *p* < 0.0001; Group: F_3,24_ = 0.5368, *p* = 0.6616; Interaction: F_15,120_ = 1.423, *p* = 0.1471) CB_2_^f/f^ and Advillin^CRE/+^; CB_2_^f/f^ mice irrespective of group; Sidak’s multiple comparisons test revealed that morphine increased mechanical paw withdrawal thresholds and decreased cold sensitivity on day 1 of injections compared to paclitaxel baseline for all groups (*p* < 0.0001). By day 6 of morphine dosing mechanical allodynia was observed in all groups, indicating the development of morphine tolerance (*p* > 0.9129 compared to paclitaxel baseline). Enhanced cold sensitivity was observed on day 6 of repeated morphine dosing compared to the post-paclitaxel baseline in all groups (*p* < 0.0369) other than the male Advillin^CRE/+^; CB_2_^f/f^ Mor+VEH group (*p* = 0.1576 difference from paclitaxel baseline), potentially indicative of opioid-induced hyperalgesia. Co-administration of morphine and LY2828360 (10 and 0.1 mg/kg per day respectively, i.p. x 6 days) in Phase II increased mechanical withdrawal thresholds (**Fig. 7B**) and decreased cold sensitivity (**Fig. 7C**) in male CB_2_^f/f^ but not male Advillin^CRE/+^; CB_2_^f/f^ mice (**Mechanical:** Time: F_5,120_ = 165.8, *p* < 0.0001; Group: F_3,24_ = 40.39, *p* < 0.0001; Interaction: F_15,120_ = 7.958, *p* < 0.0001; **Cold:** Time: F_5,120_ = 162.8, *p* < 0.0001; Group: F_3,24_ = 16.75, *p* < 0.0001; Interaction: F_15,120_ = 11.82, *p* < 0.0001). This morphine-sparing effect of LY2828360 treatment was absent in females (**Mechanical:** Time: F_5,125_ = 130, *p* < 0.0001; Group: F_3,25_ = 0.2787, *p* = 0.8403; Interaction: F_15,125_ = 2.225, *p* = 0.0086; **Cold:** Time: F_5,120_ = 208.7, *p* < 0.0001; Group: F_3,24_ = 2.456, *p* = 0.0876; Interaction: F_15,120_ = 2.591, *p* = 0.0021); Sidak’s multiple comparisons test revealed increased mechanical withdrawal threshold and decreased cold sensitivity on days 7-12 in male: Mor+LY mice (*p* < 0.0001 compared to paclitaxel baseline), but not male: Mor+VEH or female mice. By day 12, all groups showed enhanced cold sensitivity compared to paclitaxel baseline (*p* < 0.0006) with the exception of male Advillin^CRE/+^; CB_2_^f/f^ Mor+VEH group that did not differ from paclitaxel baseline (*p* = 0.0617).

**Fig. 7.**
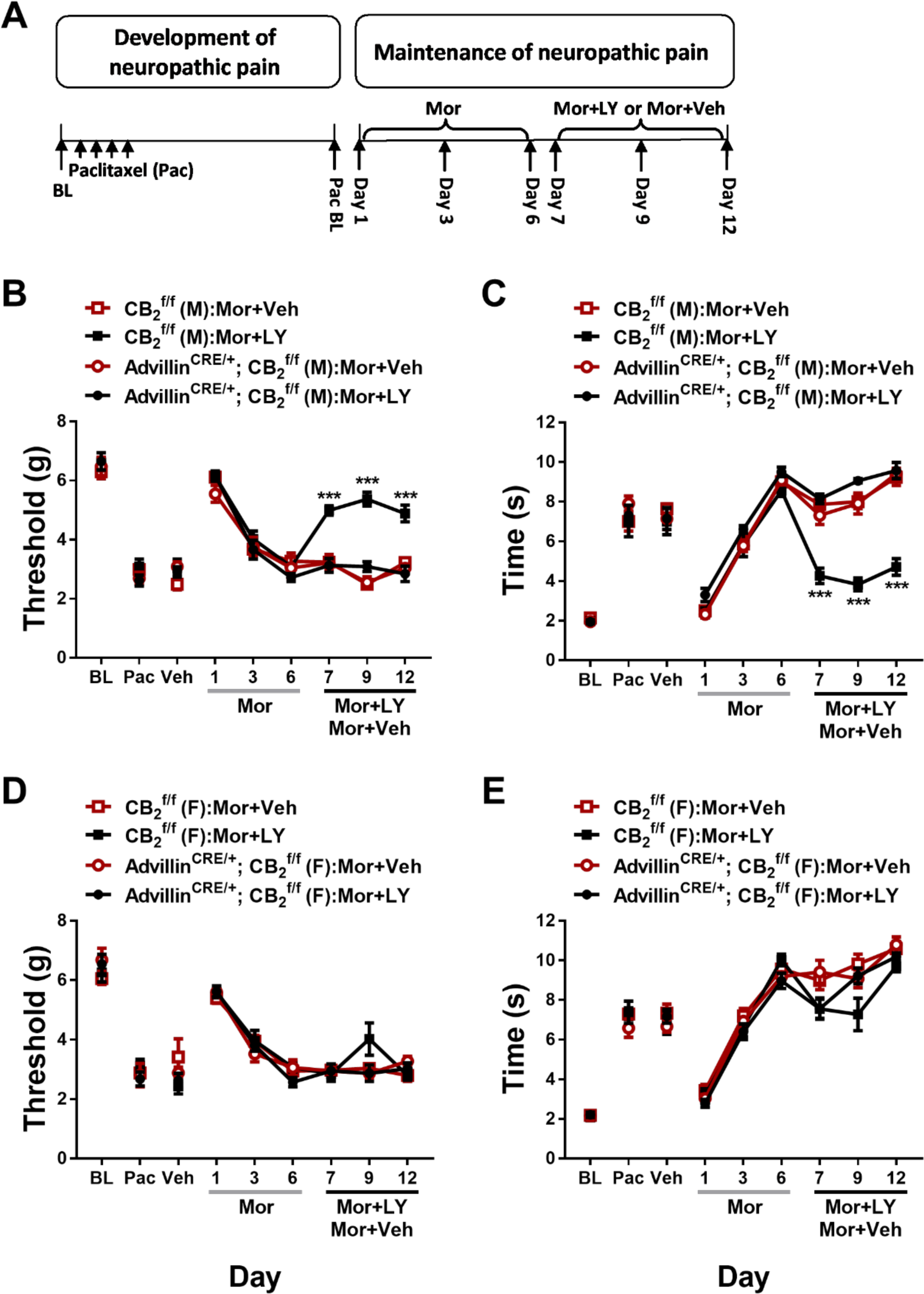
Co-administration with LY2828360 reverses morphine tolerance in male, but not female CB_2_^f/f^ mice, and had no effect on morphine tolerance in Advillin^CRE/+^; CB_2_^f/f^ mice of either sex. (A) Schematic shows the injection schedule used to evaluate the two-phase treatment during the maintenance phase of paclitaxel-induced neuropathic pain. Chronic administration of morphine (10 mg/kg per day i.p. x 6 days) initially alleviated paclitaxel-evoked mechanical (A, D) and cold (C, E) allodynia in both male and female CB_2_^f/f^ and Advillin^CRE/+^; CB_2_^f/f^ mice, but later resulted in tolerance. Co-administration of morphine with low-dose LY2828360 (0.1 mg/kg per day i.p. x 6 days) caused a reversal of tolerance to morphine’s mechanical (B) and cold (C) anti-allodynic effects in male CB_2_^f/f^, but not Advillin^CRE/+^; CB_2_^f/f^ mice. LY2828360 had no effect on mechanical (D) or cold (E) allodynia in female CB_2_^f/f^ or Advillin^CRE/+^; CB_2_^f/f^ mice and did not block the development of tolerance to the anti-allodynic effects of morphine (n = 6-8 per group). Data expressed as mean (±SEM), \*\*\**p* < 0.001.

In paclitaxel-treated CB_2_^f/f^ mice, naloxone challenge produced characteristic jumping behavior that was higher in females compared to males and withdrawal jumps were lower overall in CB_2_^f/f^ mice receiving concurrent LY2828360 + morphine compared to morphine alone (**CB_2_^f/f^:** Treatment: F_1,28_ = 4.903, *p* = 0.0351; Sex: F_1,28_ = 4.355, *p* = 0.0461; Interaction: F_1,28_ = 0.0798, *p* = 0.7796; **Fig. 8A**). By contrast, in Advillin^CRE/+^; CB_2_^f/f^ mice receiving the same treatments, no effect of sex or concurrent LY2828360 treatment was observed (**Advillin^CRE/+^; CB_2_^f/f^:** Treatment: F_1,21_ = 0.09588, *p* = 0.7599; Sex: F_1,21_ = 0.5371, *p* = 0.4717; Interaction: F_1,21_ = 0.2613, *p* = 0.6146; **Fig. 8B**).

**Fig. 8.**
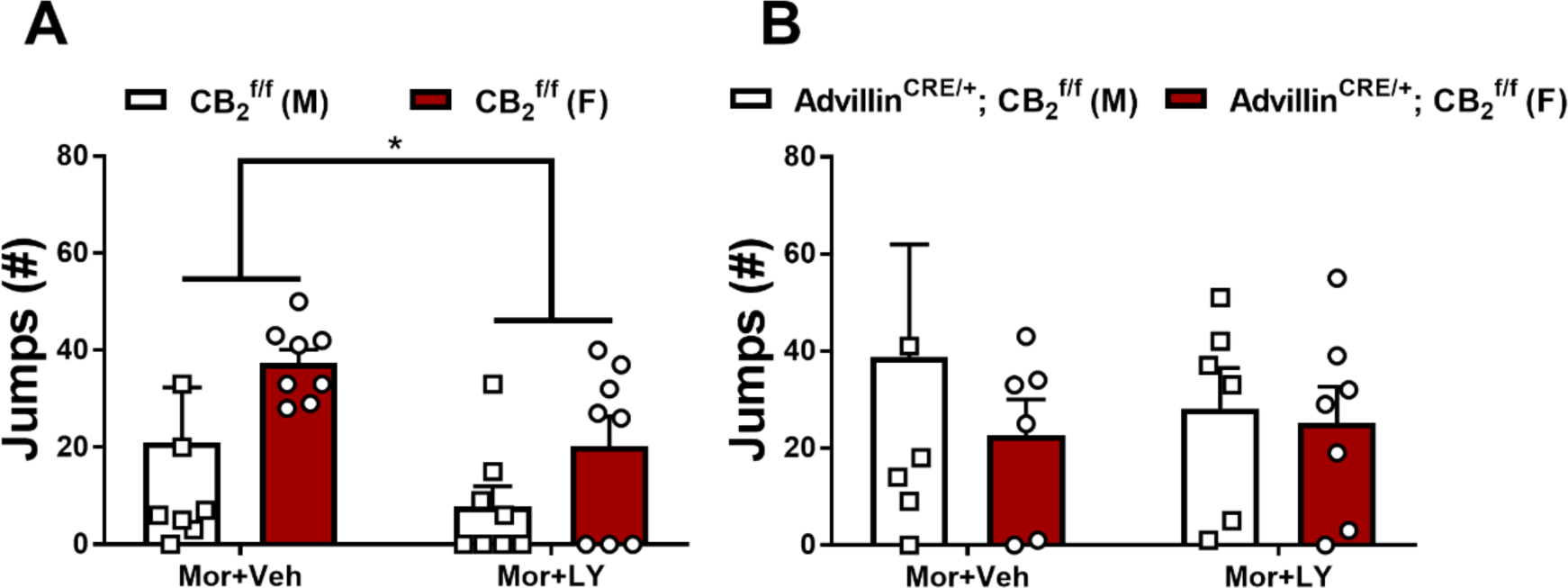
Impact of co-administration of low-dose LY2828360 with morphine (0.1 and 10 mg/kg per day i.p. x 6 days, respectively) on naloxone-precipitated morphine withdrawal in male and female CB_2_^f/f^ and Advillin^CRE/+^; CB_2_^f/f^ mice. CB_2_^f/f^ mice treated with morphine + vehicle instead of LY2828360 showed an increased number of jumps compared to the LY2828360 + morphine group. No difference in withdrawal jumps was observed between groups in Advillin^CRE/+^; CB_2_^f/f^ mice (n = 6-8 per group). Data expressed as mean (±SEM), \**p* < 0.05.

### Experiment 6. The CB_2_ agonist LY2828360 dose dependently reduced mechanical and cold allodynia in both CB_2_^f/f^ and CX3CR1^CRE/+^; CB_2_^f/f^ mice

LY2828360 dose-dependently increased mechanical paw withdrawal thresholds (**Fig. 9A**) and decreased cold sensitivity (**Fig. 9B**) irrespective of genotype in paclitaxel-treated CB_2_^f/f^ and CX3CR1^CRE/+^; CB_2_^f/f^ mice (**Mechanical:** Dose: F_4,112_ = 121.4, *p* < 0.0001; Genotype: F_1,28_ = 0.3664, *p* = 0.5499; Interaction: F_4,112_ = 0.3888, *p* = 0.8163; **Cold:** Dose: F_4,112_ = 91.56, *p* < 0.0001; Genotype: F_1,28_ = 0.7823, *p* = 0.3840; Interaction: F_4,112_ = 2.179, *p* = 0.0758). LY2828360 did not alter mechanical (**Fig. 9C**) or cold (**Fig. 9D**) responsiveness in CB_2_^f/f^ or CX3CR1^CRE/+^; CB_2_^f/f^ mice treated with cremophor-based vehicle in lieu of paclitaxel (**Mechanical:** Dose: F_4,40_ = 1.883, *p* = 0.1322; Genotype: F_1,10_ = 0.00136, *p* = 0.9713; Interaction: F_4,40_ = 1.6, *p* = 0.1931; **Cold:** Dose: F_4,40_ = 10.1, *p* < 0.0001; Genotype: F_1,10_ = 0.2299, *p* = 0.6419; Interaction: F_4,40_ = 1.016, *p* = 0.4106).

**Fig. 9.**
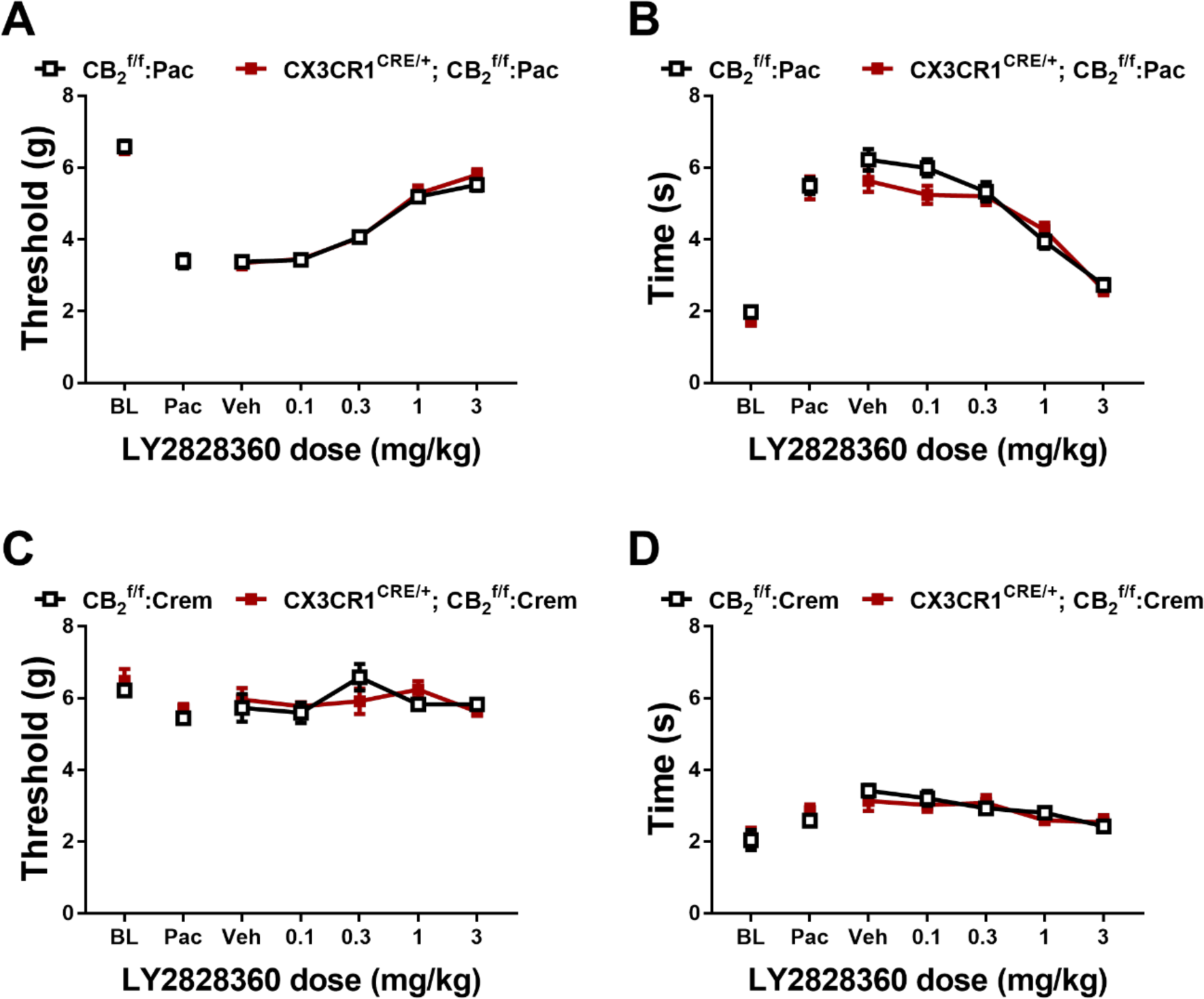
LY2828360 dose-dependently alleviated paclitaxel evoked mechanical (A) and cold (B) allodynia in both CB_2_^f/f^ and CX3CR1^CRE/+^; CB_2_^f/f^ mice. LY2828360 did not alter mechanical (C) or cold (D) sensitivity in CB_2_^f/f^ and CX3CR1^CRE/+^; CB_2_^f/f^ control mice treated with cremophor vehicle (n = 6-16 per group). Data expressed as mean (±SEM).

### Experiment 7. Previous chronic administration of LY2828360 blocked the development of morphine tolerance in male, but not female CB_2_^f/f^ and CX3CR1^CRE/+^; CB_2_^f/f^ mice

To examine the effects of LY2828360 treatment on the development of morphine tolerance in CX3CR1^CRE/+^; CB_2_^f/f^ mice, a two-phase pharmacological manipulation was used after neuropathic nociception was established and stable (**Fig. 10A**). Phase I treatment with LY2828360 (3 mg/kg per day i.p. x 12 days) increased mechanical withdrawal thresholds (**Fig. 10B,D**) and decreased cold sensitivity (**Fig. 10C,E**) in both male (**Mechanical:** Time: F_5,70_= 54.57, *p* < 0.0001; Genotype: F_1,14_ = 0.07038, *p* = 0.7946; Interaction: F_5,70_ = 0.279, *p* = 0.9232; **Cold:** Time: F_5,70_ = 87.75, *p* < 0.0001; Genotype: F_1,14_ = 0.4061, *p* = 0.5342; Interaction: F_5,70_ = 0.2365, *p* = 0.9451) and female (**Mechanical:** Time: F_5,55_ = 52.19, *p* < 0.0001; Genotype: F_1,11_ = 0.2598, *p* = 0.6203; Interaction: F_5,55_ = 0.9811, *p* = 0.4376; **Cold:** Time: F_5,55_ = 58.25, *p* < 0.0001; Genotype: F_1,11_ = 0.1961, *p* = 0.6665; Interaction: F_5,55_ = 0.6459, *p* = 0.6657) mice irrespective of genotype; Sidak’s multiple comparisons test revealed increased mechanical withdrawal threshold and decreased cold sensitivity on all phase I days tested compared to paclitaxel baseline in both sexes (*p* < 0.0001). Phase II treatment with morphine (10 mg/kg per day i.p. x 12 days) increased mechanical withdrawal thresholds (**Fig 10B,D**) and decreased cold sensitivity (**Fig. 10C,E**) without the development of tolerance in male mice (**Mechanical:** Time: F_6,84_ = 102.8, *p* < 0.0001; Genotype: F_1,14_ = 0.1651, *p* = 0.6907; Interaction: F_6,84_ = 0.4404, *p* = 0.8498; **Cold:** Time: F_6,84_ = 111.3, *p* < 0.0001; Genotype: F_1,14_ = 3.658, *p* = 0.0765; Interaction: F_6,84_ = 0.9566, *p* = 0.4597) but this effect was absent in female mice (**Mechanical:** Time: F_6,66_ = 65.94, *p* < 0.0001; Genotype: F_1,14_ = 0.719, *p* = 0.4146; Interaction: F_6,66_ = 1.327, *p* = 0.2577; **Cold:** Time: F_6,66_ = 81.02, *p* < 0.0001; Genotype: F_1,14_ = 0.09744, *p* = 0.7608; Interaction: F_6,66_ = 1.36, *p* = 0.2437); Sidak’s multiple comparisons test revealed that acute treatment with morphine in Phase II (day 16) increased mechanical paw withdrawal thresholds and decreased cold sensitivity compared to day 15 post-paclitaxel baseline responding in both sexes regardless of genotype (*p* < 0.0001 for all comparisons). Morphine tolerance developed in female mice of both genotypes at subsequent time points (*p* < 0.0218 on days 19-27 for all timepoints compared to day 16), whereas anti-allodynic effects of morphine were preserved in male mice of both genotypes for the duration of phase 2 (*p* > 0.0233 on days 19-27 compared to day 16). In paclitaxel treated CB_2_^f/f^ and CX3CR1^CRE/+^; CB_2_^f/f^ mice with a history of chronic treatment with LY282860 and morphine, naloxone challenge produced characteristic jumping behavior that did not differ as a function of genotype or sex (Sex: F_1,24_ = 0.6948, *p* = 0.4127; Genotype: F_1,24_ = 0.5811, *p* = 0.4533; Interaction: F_1,24_ = 1.904, *p* = 0.1804; **Fig. 11**).

**Fig. 10.**
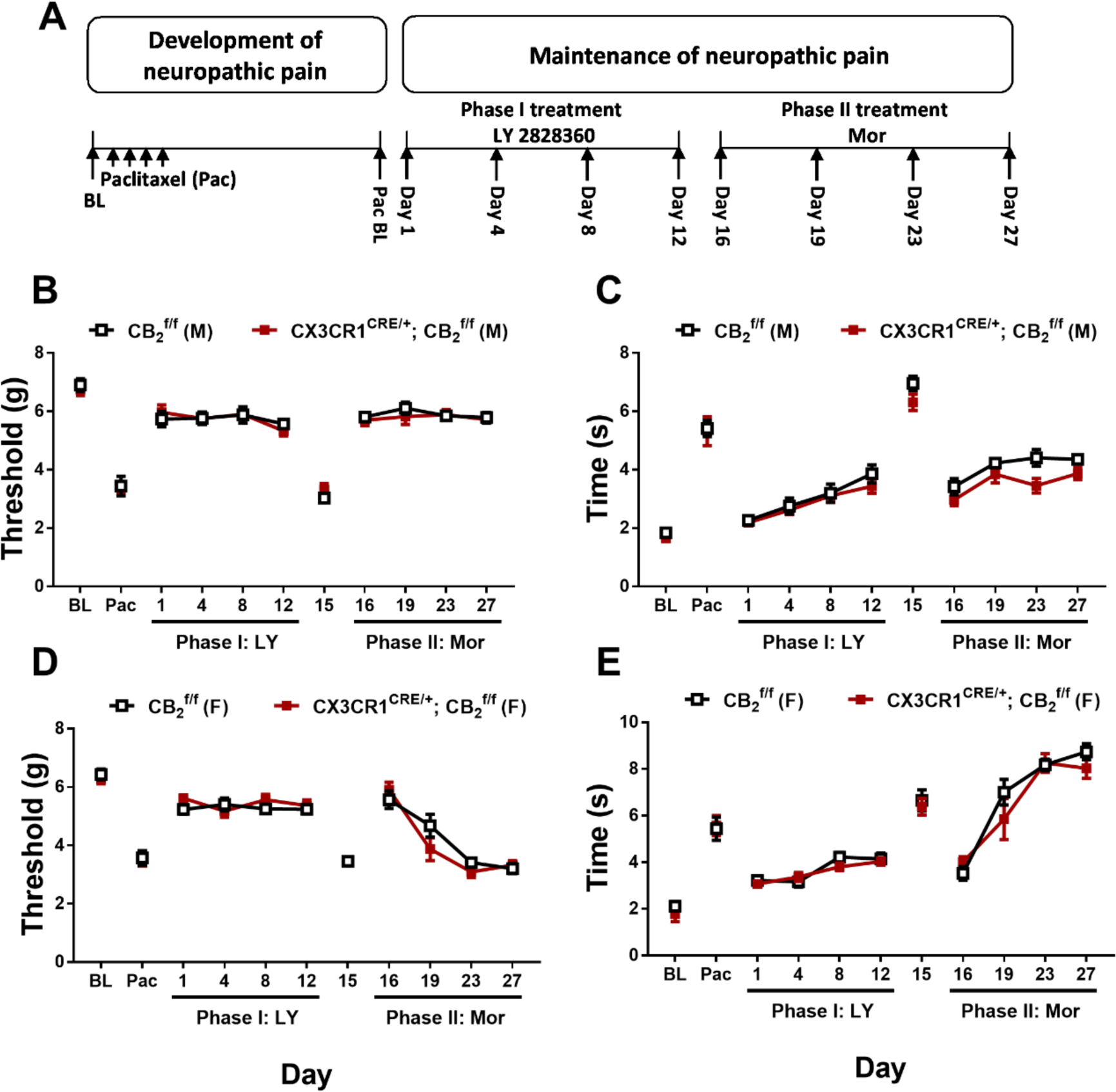
History of chronic LY2828360 treatment blocked the development of morphine tolerance in male, but not female CB_2_^f/f^ and CX3CR1^CRE/+^; CB_2_^f/f^ mice. (A) Schematic shows the injection schedule used to evaluate the two-phase treatment during the maintenance phase of paclitaxel-induced neuropathic pain. Chronic LY2828360 treatment (3 mg/kg per day i.p. x 12 days) alleviated paclitaxel evoked mechanical (B, D) and cold (C, E) allodynia and blocked the development of tolerance to the anti-allodynic effects of morphine (10 mg/kg per day i.p. x 12 days) in male (B, C), but not female (D, E) CB_2_^f/f^ and CX3CR1^CRE/+^; CB_2_^f/f^ mice (n = 6-8 per group). Data expressed as mean (±SEM).

**Fig. 11.**
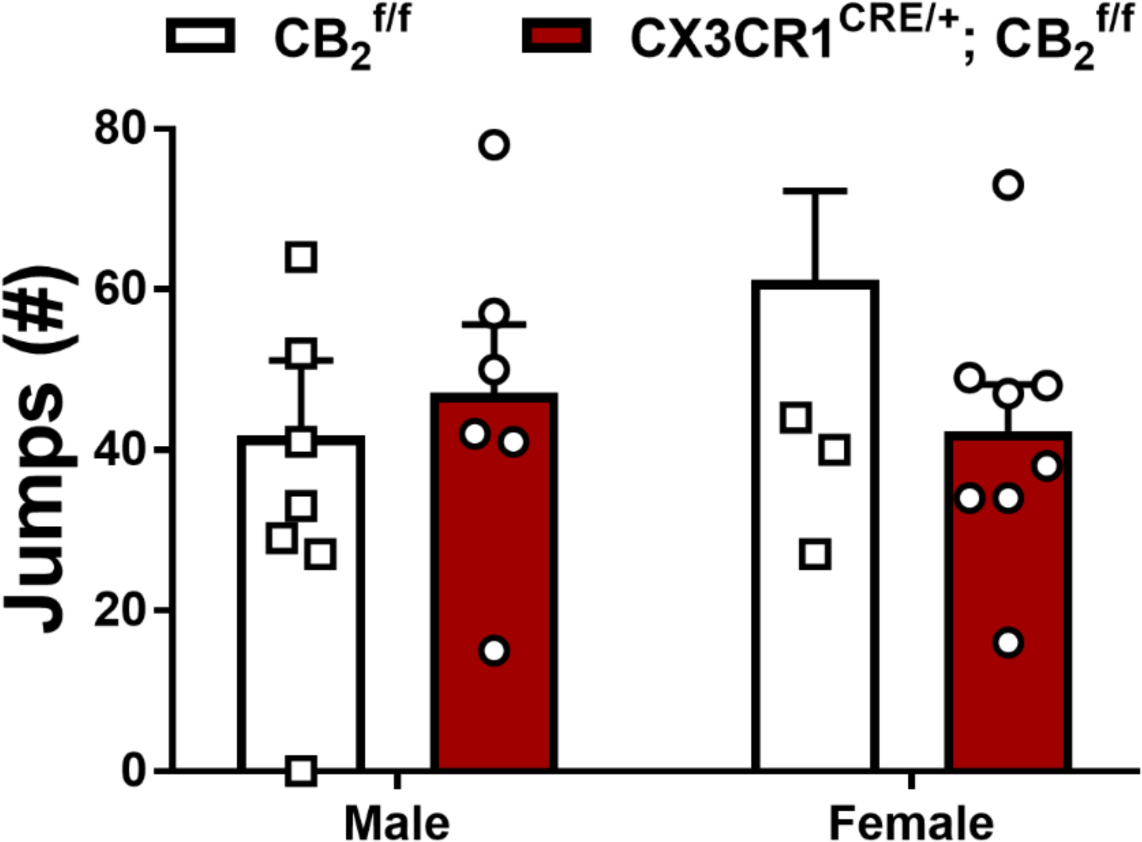
Impact of a history of chronic LY2828360 (3 mg/kg per day i.p. x 12 days) on naloxone-precipitated morphine withdrawal in male and female CB_2_^f/f^ and CX3CR1^CRE/+^; CB_2_^f/f^ mice (n = 6-8 per group). No differences were found between genotype or sex. Data expressed as mean (±SEM), \**p* < 0.05.

### Experiment 8. Prophylactic administration of LY2828360 reduced but did not permanently prevent development of mechanical and cold allodynia in both CB_2_^f/f^ and CX3CR1^CRE/+^; CB_2_^f/f^ mice

Prophylactic treatment with LY2828360 for 7 days increased mechanical paw withdrawal thresholds and decreased cold sensitivity during the period of chronic dosing but did not permanently prevent the development of mechanical or cold allodynia in either CB_2_^f/f^ or CX3CR1^CRE/+^; CB_2_^f/f^ mice (**Mechanical:** Time: F_5,80_ = 195.1, *p* < 0.0001; Group: F_3,16_ = 19.72, *p* < 0.0001; Interaction: F_15,80_ = 10.11, *p* < 0.0001; **Cold:** Time: F_5,80_ = 218.7, *p* < 0.0001; Group: F_3,16_ = 10.38, *p* = 0.0005; Interaction: F_15,80_ = 8.859, *p* < 0.0001); Sidak’s multiple comparisons test revealed that LY2828360 treatment increased paw withdrawal thresholds in all groups on day 7 of injections, regardless of genotype (*p* < 0.0001), and this effect persisted on day 11 of injections in CB_2_^f/f^ mice (*p* = 0.0050). By contrast, no difference in mechanical paw withdrawal threshold was observed between groups at other time points (*p* > 0.0840), suggesting that LY2828360 delayed, but did not prevent, the development of paclitaxel-induced mechanical allodynia. Similarly, LY2828360 decreased paclitaxel-induced cold responsiveness on day 7 and 11 following initiation of paclitaxel dosing irrespective of genotype. Thus, anti-allodynic efficacy LY282830 outlasted the period of LY2828360 treatment but eventually diminished. Decreased cold sensitivity was observed in mice receiving LY2828360 compared to Vehicle on day 7 and 11 regardless of genotype (*p* < 0.0051), and this effect still observed on day 18 in CB_2_^f/f^ mice (*p* = 0.0167), but not at other timepoints (*p* > 0.2940; **Fig. 12**).

**Fig. 12.**
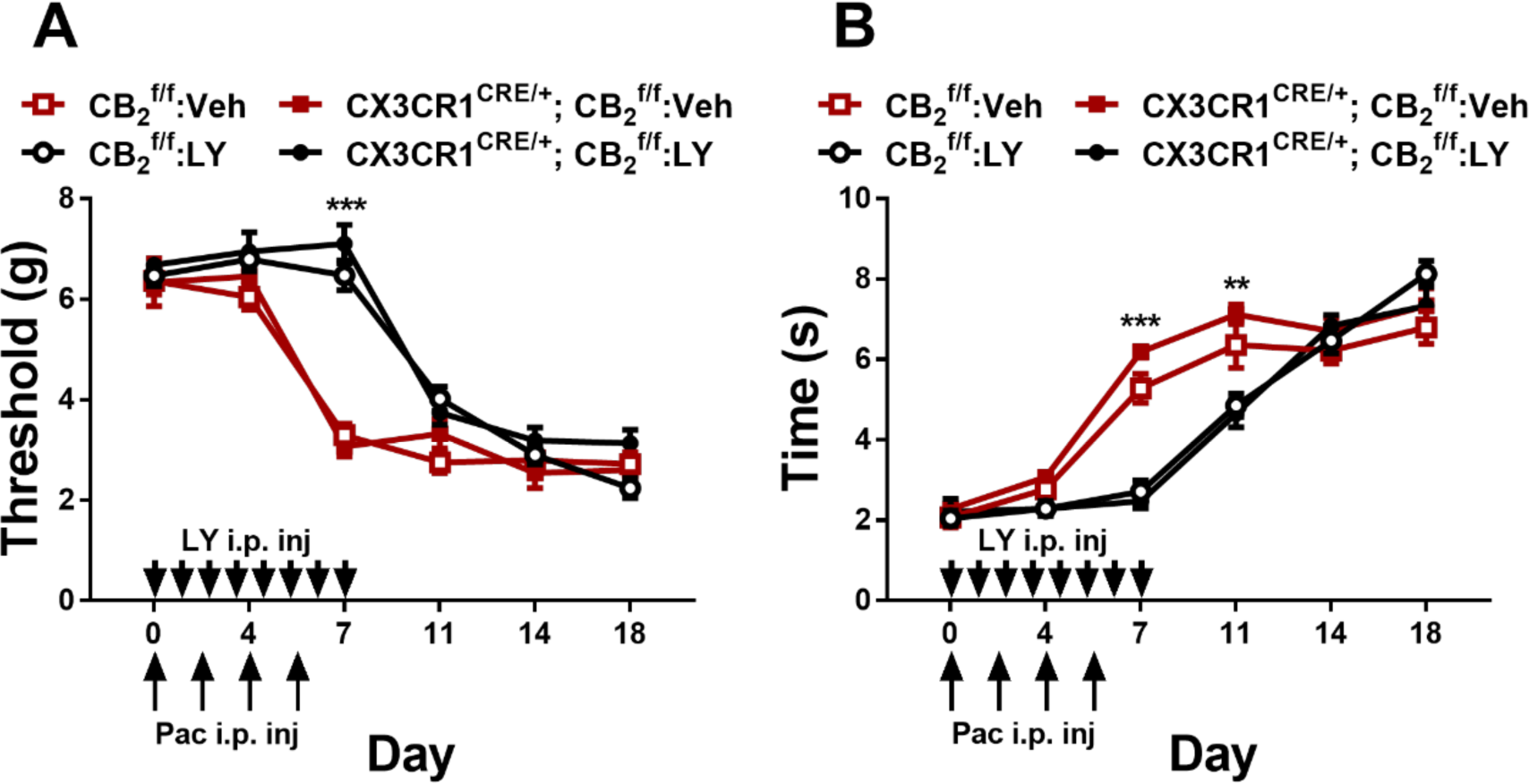
LY2828360 (3 mg/kg per day i.p. x 8 days) delayed the onset of paclitaxel-induced mechanical (A) and cold (B) allodynia but did not prevent their development. Anti-allodynic effects of LY2828360 were no longer observed 4-7 days following termination of dosing (n = 5 per group). Data expressed as mean (±SEM), \*\**p* < 0.005, \*\*\**p* < 0.001.

## Discussion

Activation of cannabinoid CB_2_ receptors suppresses certain types of pathological pain without producing unwanted side effects (e.g. Guindon and Hohmann 2008; Murineddu et al. 2013; Cabañero et al. 2021). Our lab has previously examined the ability of the slowly-signaling, G protein-biased CB_2_ receptor agonist LY2828360 to reduce neuropathic nociception in the paclitaxel model of chemotherapy-induced neuropathic nociception (Lin et. 2018; Lin et al. 2022). Here we generated mice with a conditional deletion of CB_2_ receptors from peripheral sensory neurons (Advillin^CRE/+^; CB_2_^f/f^) or from CX3CR1-expressing microglia/macrophages (CX3CR1^CRE/+^; CB_2_^f/f^) to determine the specific population of cells involved in the anti-allodynic effect of LY2828360. First, we validated our Advillin^CRE/+^; CB_2_^f/f^ mouse line using qRT-PCR and showed that both GFP and CB_2_ mRNA were reduced in the dorsal root ganglia (DRG) derived from Advillin^CRE/+^; CB_2_^f/f^ mice compared to CB_2_^f/f^ controls, whereas no difference in CB_2_ mRNA levels were observed in the spinal cord, paw skin or spleen. This finding replicates our previous work showing that Advillin^CRE/+^; CB_2_^f/f^ mice have reduced CB_2_ and GFP mRNA levels only in the DRG compared to these other tissues (Carey et al. 2023) and documents that the cKO generated did not result in global spontaneous (germline) deletions. The CX3CR1^CRE/+^; CB_2_^f/f^ mouse line used here has been successfully used by our groups to uncover behavioral phenotypes mediated by CB_2_ agonists in other studies (Behlke et al. 2022) and other models (unpublished).

We used a paclitaxel-induced neuropathy model to compare the anti-allodynic efficacy of LY2828360, which is brain penetrant, with that of AM1710, a CB_2_ agonist with limited ability to penetrate the CNS (Hollinshead et al. 2013; Rahn et al. 2011) in paclitaxel-treated CB_2_^f/f^ mice and two different cKO mouse lines (i.e. Advillin^CRE/+^; CB_2_^f/f^ mice and CX3CR1^CRE/+^; CB_2_^f/f^ mice). Both CB_2_ agonists, LY2828360 and AM1710, dose-dependently reduced mechanical and cold hypersensitivity in CB_2_^f/f^ mice but lacked anti-allodynic efficacy in Advillin^CRE/+^; CB_2_^f/f^ mice. We also observed anti-allodynic efficacy of LY2828360 in CX3CR1^CRE/+^; CB_2_^f/f^ mice. These observations support our hypothesis that CB_2_ receptors in peripheral sensory neurons are necessary for the anti-allodynic effects of both LY2828360 and AM1710 in paclitaxel-induced peripheral neuropathy. By contrast, CB_2_ receptors found on CX3CR1 expressing microglia/macrophages do not appear to be involved in the anti-allodynic efficacy of these CB_2_ agonists. A similar effect was observed in our prior study showing that LY2828360 retains efficacy in CX3CR1^CRE/+^; CB_2_^f/f^ but not Advillin^CRE/+^; CB_2_^f/f^ mice in a carrageenan model of inflammatory pain (Guenther et al. 2023). Additionally, LY2828360 reduced neuropathic nociception produced by the antiretroviral agent 2,3-dideoxycytidine (ddC) in CB_2_^f/f^ but not in Advillin^CRE/+^; CB_2_^f/f^ mice (Carey et al. 2023). Together these studies demonstrate an important role for CB_2_ receptors in peripheral sensory neurons in CB2-mediated analgesic efficacy in various pain subtypes. By contrast, CB_2_ receptors in CX3CR1 expressing microglia/macrophages are unlikely to be involved in this phenomenon. Our CX3CR1^CRE/+^; CB_2_^f/f^ mouse model is specific to microglia expressing the C-X3-C Motif Chemokine Receptor 1 gene; therefore our results do not preclude the possibility that other microglia subtypes may contribute to the anti-allodynic effect of CB_2_ agonists in our studies of inflammatory nociception (Guenther et al. 2023) and toxic neuropathy (Carey et al. 2023) or in other pain models.

Tolerance limits the therapeutic potential of analgesics such as morphine for treating chronic pain (Rosenblum et al. 2008). Our lab has previously shown sustained anti-allodynic efficacy of LY2828360 in the paclitaxel model after repeated dosing in mice (Lin et al. 2018; Carey et al. 2023). We also showed that prior treatment with LY2828360 blocked the development of tolerance to morphine in paclitaxel-treated male mice (Lin et al. 2018), however the cell types underlying this effect were unknown prior to the present report. Here we examined the ability of LY2828360 to prevent or reverse morphine tolerance in CB_2_^f/f^ and Advillin^CRE/+^; CB_2_^f/f^ mice. In paclitaxel-treated male mice, LY2828360 blocked the development of morphine tolerance; a history of 12-day chronic LY2828360 treatment (3 mg/kg i.p.) prevented subsequent development of tolerance to the anti-allodynic effects of morphine (10 mg/kg i.p. x 12 days) in male CB_2_^f/f^ and male CX3CR1^CRE/+^; CB_2_^f/f^ mice. This effect was sex dependent, as tolerance to the anti-allodynic effects of morphine developed normally in female mice after 3 days in both genotypes, indicating that CB_2_ receptors only play a critical role in preventing the development of morphine tolerance in male mice. LY2828360 also failed to prevent tolerance to the anti-allodynic effects of morphine in Advillin^CRE/+^; CB_2_^f/f^ mice in either sex, presumably because only male, but not female, mice show the morphine tolerance-sparing effect of LY2828360. These results suggest that CB_2_ receptors in peripheral sensory neurons but not CX3CR1 expressing microglia/macrophages are necessary for preventing the development of tolerance to the anti-allodynic effects of morphine and that this phenomenon is specific to male mice only. To measure morphine withdrawal in these mice, we challenged mice with naloxone at the end of the 12-day chronic morphine period and measured jumping behavior as a readout of naloxone-precipitated opioid withdrawal. We found that both male CB_2_^f/f^ and male Advillin^CRE/+^; CB_2_^f/f^ mice showed less jumping behavior in response to naloxone compared to females of both genotypes,whereas no sex differences were observed when comparing CB_2_^f/f^ mice to CX3CR1^CRE/+^; CB_2_^f/f^ mice. Thus, CB_2_ receptors in peripheral sensory neurons do not appear to be involved in naloxone-precipitated opioid withdrawal behavior as there was no difference in number of withdrawal jumps between male CB_2_^f/f^ and Advillin^CRE/+^; CB_2_^f/f^ mice. Notably, we observed genotypic differences in the development of tolerance to the anti-allodynic effects of morphine in the same mice that received chronic morphine treatment following chronic treatment with LY2828360, with sparing of morphine tolerance observed in male CB_2_^f/f^ but not male Advillin^CRE/+^; CB_2_^f/f^ mice.

Interestingly, LY2828360 was also able to reverse established morphine tolerance in paclitaxel-treated mice. Tolerance developed in both CB_2_^f/f^ and Advillin^CRE/+^; CB_2_^f/f^ mice after 3 days of repeated morphine administration and persisted for the entire morphine administration period. When a low dose of LY2828360 (0.1 mg/kg i.p.) was co-administered with morphine after the establishment of morphine tolerance, we observed a restoration of morphine antinociceptive efficacy, consistent with reversal of tolerance to morphine’s anti-allodynic effects. As was seen in the previous morphine tolerance experiment (i.e. in which administration of LY2828360 and morphine never overlapped), this effect was sex dependent as tolerance to morphine’s anti-allodynic effect was maintained in female mice even after co-administration of LY2828360 began. These observations suggest that CB_2_ receptor activation reduces morphine tolerance in the paclitaxel model in male mice only, a finding which may have clinical implications. Additionally, this reversal of morphine tolerance was not observed in Advillin^CRE/+^; CB_2_^f/f^ mice in either sex, highlighting the importance of CB_2_ receptors in peripheral sensory neurons in the maintenance of tolerance to morphine’s anti-allodynic effects in males, the only sex in which LY2828360-induced sparing of morphine tolerance was observed (but see Carey et al. 2023).

We also compared naloxone-precipitated morphine withdrawal in CB_2_^f/f^ and Advillin^CRE/+^; CB_2_^f/f^ mice of both sexes. We observed fewer withdrawal jumps in CB_2_^f/f^ mice that received morphine co-administered with LY2828360 compared to vehicle. These observations suggest that a behaviorally inactive dose of LY2828360, when co-administered with morphine, was sufficient to attenuate withdrawal behavior. By contrast, no protective effect of LY2828360 was observed on withdrawal behavior under the conditions of the two-phase study because LY2828360 was never given concurrently with morphine. In that two-phase study, dosing with LY2828360 terminated 4 days prior to initiation of morphine dosing, and drug effects were likely washed out by the time withdrawal was precipitated with naloxone. By contrast, LY2828360 was on board when co-administered with morphine and was able to impact withdrawal behavior following naloxone challenge, and sex differences in the withdrawal phenotype were not observed. Lastly, in morphine-dependent Advillin^CRE/+^; CB_2_^f/f^ mice we did not observe sex differences or impact of LY2828360 treatment following naloxone-precipitated withdrawal in this experiment.

We also investigated whether chronic prophylactic administration of LY2828360 (3 mg/kg i.p.) for 7 days could prevent the development of peripheral neuropathic nociception during the establishment of paclitaxel-induced peripheral neuropathy. LY2828360 delayed, but did not prevent, the development of paclitaxel-induced mechanical and cold allodynia in both CB_2_^f/f^ and CX3CR1^CRE/+^; CB_2_^f/f^ mice. Mechanical and cold hypersensitivity returned following termination of dosing with LY2828360.

Elucidation of therapeutic strategies to prevent or block morphine tolerance has been the focus of significant research efforts (Habibi-Asl et al. 2014; Mansouri et a. 2015; Hassanipour et al. 2016; Hosseinzadeh et al. 2016). The present results show that the CB_2_ agonist LY2828360 can enhance the anti-allodynic effects of morphine by both preventing and reversing morphine tolerance. However, much of the literature, including our previous report (Lin et al. 2018) assessed morphine tolerance in male subjects only. The present study replicates findings from Lin et al. (2018) showing that chronic pre-treatment with the CB_2_ agonist LY2828360 can prevent the development of morphine tolerance in a mouse model of paclitaxel-induced neuropathic pain and shows that this phenomenon is preferentially observed in male, but not female, mice. This report also expands upon our initial demonstration of this phenomenon by establishing the cell types that express CB_2_ that are responsible for this phenomenon. We show that peripheral sensory neuron CB_2_ receptors are required for the anti-allodynic effects of CB_2_ agonists in this model. We also show that activation of the CB_2_ receptor can reverse already established morphine tolerance in mice. These results are significant given that many human patients may have pre-existing tolerance to opioids, and, consequently, treatment with LY2828360 may be beneficial in reinstating the anti-allodynic effects of morphine. Overall, LY2828360 may be a useful first line treatment for chemotherapy-induced peripheral neuropathy, as it can supress pathological pain with sustained efficacy, and both prevent and reverse morphine tolerance. Whereas CB_2_ agonists were effective in suppressing neuropathic nociception in mice of both studies, the present results suggest that this is not the case for morphine sparing effects, in which a notable sexual dimorphism in sparing of morphine tolerance was observed. Our studies also suggest that a previously underappreciated role of sex may be an important factor to consider in any clinical studies of CB_2_-opioid interactions.

## Acknowledgments

Supported by DA047858 (to AGH and KM), DA009158 (to AM and AGH), an Indiana Addiction Grand Challenge Grant (to AGH), the Research and Education Component of the Advancing a Healthier Wisconsin Endowment at the Medical College of Wisconsin (to CH), and Ministerio de Ciencia e Innovación – Agencia Estatal de Investigación and Fondo Europeo de Desarrollo Regional (Proyecto PID2022-138461OB-I00, supported by MCIN /AEI /10.13039/501100011033 / FEDER, UE (to JR). KG is a Harlan Research Scholar.

